# Tethered Bichromophoric Fluorophore Quencher Voltage Sensitive Dyes

**DOI:** 10.1101/473843

**Authors:** Ping Yan, Corey Acker, Leslie M. Loew

## Abstract

Voltage sensitive dyes (VSDs) are used for *in vitro* drug screening and for *in vivo* imaging of patterns of electrical activity. However, wide application of this technology is limited by poor sensitivity. A promising approach uses a 2-component system consisting of charged membrane permeable quenchers together with fluorophores labeling one side of the membrane; this produces voltage-dependent fluorescence quenching. However, to achieve good sensitivity, the quencher compound must be used at high concentrations, which can perturb the membrane capacitance or have other pharmacological effects. By developing tethered bichromophoric fluorophore quencher (TBFQ) dyes, where the fluorophore and quencher are covalently connected by a long hydrophobic chain, the concentration required is minimized, and the sensitivity is maximized. A series of 13 TBFQ dyes based on the AminoNaphthylEthenylPyridinium (ANEP) fluorophore and the dipicrylamine anion (DPA) quencher have been synthesized and tested in an artificial lipid bilayer apparatus. The best one from the screening, **TBFQ1**, shows a 2.5 fold change in fluorescence per 100mV change in membrane potential, and the response kinetics is in 10-20 ms range. This sensitivity is an order of magnitude better than commonly used fluorescent voltage sensors. The design principles for TBFQ VSDs described here can be readily extended to other spectral regions and promise to greatly enhance our ability to monitor electrical activity in cells and tissues.

## Introduction

Electrical signaling is the primary means of communication between neurons and constitutes the fundamental mechanism for information processing in the brain. Electrical activity also controls the contraction of muscle and, in particular, the precise propagation of action potentials is the primary means for control of heart rhythm. Electrode-based intracellular recordings provide the highest fidelity means for measuring the fast electrical signals, but it is impractical to record from more than a few cells in this way – especially *in vivo*. On the other hand, imaging approaches, usually with fluorescent sensors, have the advantage that patterns of activity can be studied with high resolution over large areas of brain or heart by employing sensitive high frame-rate imaging cameras.^1^ Calcium imaging is the most common technology because fluorescent calcium sensors produce large changes in emission upon binding calcium.^2^ However, calcium changes are not a direct correlate of electrical activity and, in particular, do not properly reflect the shape of action potentials, oscillations, spiking, inhibitory signals or subthreshold potentials. A more direct analysis of electrical activity is enabled through imaging of fluorescence from molecular voltage sensors which include both voltage-sensitive organic dyes (VSDs)^3^ and genetically encoded fluorescent protein based voltages indicators (GEVIs).^4^ However, fluorescence voltage sensing currently has severe sensitivity limitations: sensitivity is, at best, ~30% change in fluorescence per action potential on single cells; in brain experiments, where background sensor fluorescence from large populations of labeled but inactive cells will dilute the optical signal, the sensitivity can be orders of magnitude lower. Thus, signal to noise ratio can be a severely limiting factor for voltage imaging applications that require high spatial and temporal resolution.

To follow action potentials, VSDs need to respond to the voltage change across the membrane in milliseconds or faster. This need for speed is what places limits on the range of mechanisms for the design of VSDs. Our lab has focused on the use of a molecular Stark effect (also called electrochromism), whereby an electron rearrangement between the ground and excited states causes an electric field-dependent shift in the VSD spectrum.^5^ Recently, an electron transfer mechanism has been developed as a basis for a new class of VSDs.^3d, 6^ Because these mechanisms are fundamentally electronic, they are guaranteed to be fast. However, because these mechanisms sense the voltage drop only across the chromophore of the VSD, most of the voltage drop across the membrane is going to waste. Therefore, if a VSD could sample the entire thickness of the membrane, it might be a more sensitive transducer of the full membrane potential.

In the mid ‘90s, Roger Tsien’s lab developed two-component voltage sensors composed of Förster resonance energy transfer (FRET) donor-acceptor pairs.^3e, 7^ The idea is that a cationic acceptor fluorophore is anchored to the external surface of the cell and a permeable anion donor fluorophore moves from the outer to the inner surface in response to depolarization; depolarization thereby effects an increase in donor fluorescence and a decrease in acceptor fluorescence. Sensitivity is improved because the schemes utilize the entire voltage drop across the membrane. This approach has been successfully applied in high throughput cell based drug screening assays, but was not found very practical for voltage imaging. This is because the redistribution of dyes was often slow in tissue and optimal sensitivity could require very high, potentially toxic, concentrations of the sensor molecules. A variant on this approach was developed by the Bezanilla and DiGregorio labs, who used dipicrylamine (DPA) as a general purpose permeable anionic quencher.^8^ Here, almost any fluorophore that could be anchored to the external surface of the cell membrane could be quenched by DPA anion in a voltage-dependent manner. Additionally, DPA itself is non-fluorescent. This system could be used to image electrical activity in brain slices with sensitivities of ~40% fluorescent increase per action potential.^8d^ A high concentration of the DPA molecules is required to assure that at resting membrane potential they are on average sufficiently close to the externally bound fluorophore molecules to produce substantial quenching; this quenching is then relieved when the DPA anions redistribute to the interior surface upon depolarization. However the high levels of DPA required for these sensitivities could increase membrane capacitance and produce other toxic side effects.

We reasoned that covalently tethering an impermeable fluorophore to a membrane-permeable quencher with a membrane spanning hydrophobic linker could overcome the need for high sensor concentrations and could also significantly improve sensitivity. The idea, as depicted in Fig. 1, is that in one polarization state, the two chromophores would be held apart, separated by the ~4 nm thickness of the membrane. In this state, the sensor would emit fluorescence upon excitation. In the opposite polarization state, the quencher and fluorophore would be on the same side of the membrane, producing a dark state. In this work, we present our first approaches and initial successes toward the design and synthesis of tethered bichromophoric fluorophore quencher (TBFQ) VSDs. Because the DPA anion was proven to rapidly traverse the thickness of cell membranes in a voltage-dependent manner, we used this chromophore as the quencher moiety. We tried various fluorophores, sidechains and tethers to learn how structural features of the TBFQ system could affect sensitivity and kinetics. Our best TBFQ VSDs show response times in the ms time scale. Remarkably, we have achieved sensitivities of 2.5 fold changes in fluorescence per 100mV change in membrane potential. All previous fast voltage sensors display sensitivities of <50%/100mV.^3-4^

**Figure 1.**
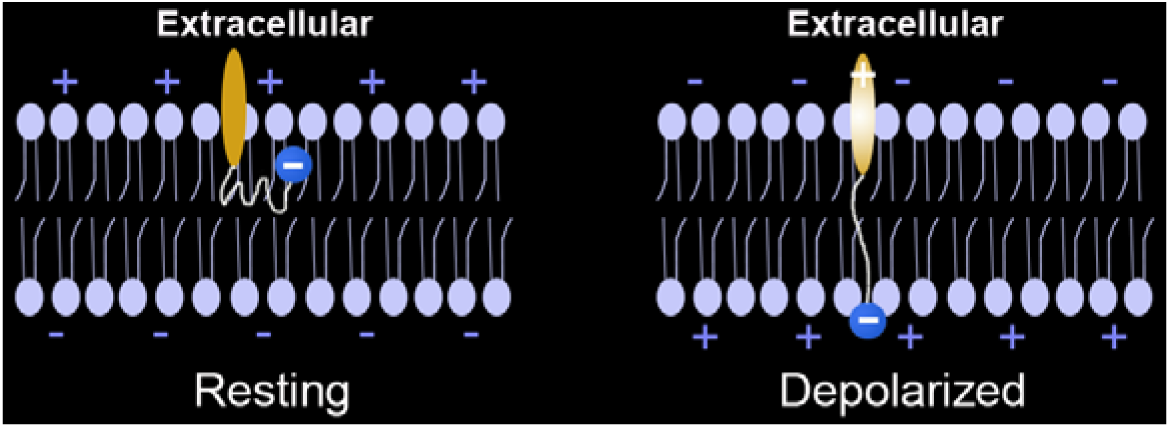
Illustration of a tethered TBFQ VSD responding to membrane potential changes. The fluorophore is anchored to the extracellular surface, and the quencher is a membrane permeable anion.

## Results and Discussion

### TBFQ design based on Förster resonance energy transfer (FRET)

FRET has been generally considered as the mechanism behind quenching by DPA anion of the fluorescence from Di-8-ANEPPS,^9^eGFP^8a^ and Di-O^8c^ in previous applications of the 2-component scheme for sensitive voltage sensing. The best reported sensitivity was for the Di-O/DPA pair, with a ΔF/F of 56%/100mV under optimal conditions.^8c^ To establish a baseline for the behavior of a 2-component system with which to compare our new TBFQ VSDs, we used Di-4-ANEPPS (which has the same hemicyanine chromophore as di-8ANEPPS) as the donor and DPA as acceptor. Di-4-ANEPPS also serves as a standard single component VSD for our comparisons as it is arguably the most commonly used fast VSD. We calculated *R*_0_ for the Di-4-ANEPPS and DPA pair to be 2.6 nm.^10^ This number is counterintuitively large considering the poor overlap between Di-4-ANEPPS emission and DPA absorbance spectra (Fig. 2). This is because the 1/6 root dependence makes the *R*_0_ somewhat insensitive to the spectral overlap. Therefore, the quenching effect should be close to 100% when the two chromophores are in close proximity on the same side of the membrane. However, this highly non-linear dependence of FRET on distance also makes the quenching effect quite small, ~7%, when the donor and acceptor are separated by the thickness of a membrane of ~4nm.

**Figure 2.**
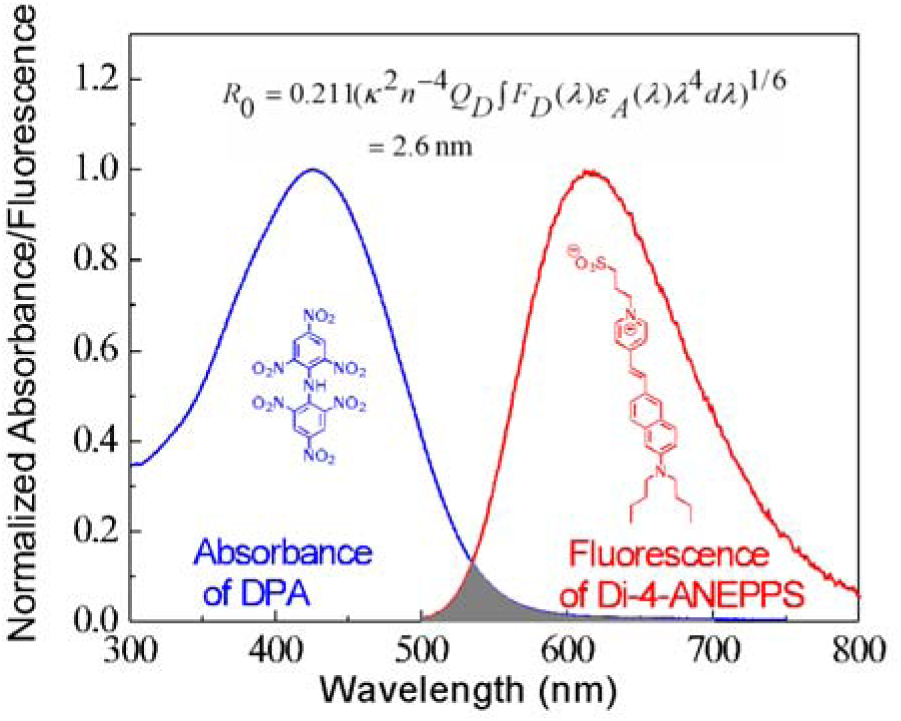
Overlap between Di-4-ANEPPS emission and DPA absorbance spectra, and *R*_0_ calculation. In the equation, *k*^2^ is the orientation factor between donor and acceptor molecules, *n* is the index of refraction, *Q_D_* is the quantum yield of donor fluorescence in the absence of acceptor, *F_D_* is the peak-normalized fluorescence spectrum of the donor, and ε_A_ is the absorption spectrum of the acceptor. The protonated structure of DPA is shown here, but it is anionic at neutral pH.

These theoretical considerations were corroborated by experiments on the D-4-ANEPPS/DPA donor/acceptor FRET pair. We used a hemispherical bilayer apparatus to apply well controlled voltage steps while recording the fluorescence from a thinly illuminated arc at the bottom of the membrane. This apparatus, which is an updated version of a system that has been previously used in this lab,^5a, 11^ is described in the Supporting Information. It consists of a lipid bilayer membrane that is expanded by hydrostatic pressure at the tip of a Teflon pipet containing 100 mM KCl. The resulting bubble is suspended in a cuvet for fluorescence recording; electrodes in the cuvet and inside the pipet serve to apply voltage steps. A comparison of the responses of Di-4-ANEPPS to the Di-4-ANEPPS/DPA pair is shown in Figure 3. The results show that wavelength dependence of the Di-4-ANEPPS in Fig. 3A is biphasic, as is characteristic of electrochromic VSDs;^5a^ these results actually represent combined effects from the voltage-dependence arising from the shifted excitation and emission spectrum, which is being detected at the red edge. In Fig. 3B, the relative fluorescence response to the voltage-dependent translocation of the DPA quencher would not be expected to be wavelength dependent; however the results display wavelength dependence because of the superposition of the electrochromic mechanism with the quencher translocation mechanism. Di-4-ANEPPS gives an approximately linear voltage dependence (Fig. 3C) over a broad range of applied voltage, but with a modest sensitivity of 12%/100mV, as reported previously.^11b^ The donor/acceptor pair produce a much larger peak response of 16%/100mV in the range of - 50 mV to +50 mV. The latter is non-linear, as would be expected from a voltage-dependent redistribution of DPA across the membrane; it also depends on the initial placement of the donor and acceptor pairs on opposite sides of the membrane; if they are added to the same side, we are unable to observe any voltage sensitivity. The differences in the behavior displayed in Fig. 3 allow us to distinguish between a pure electrochromic VSD response and the quencher translocation mechanism.

**Figure 3.**
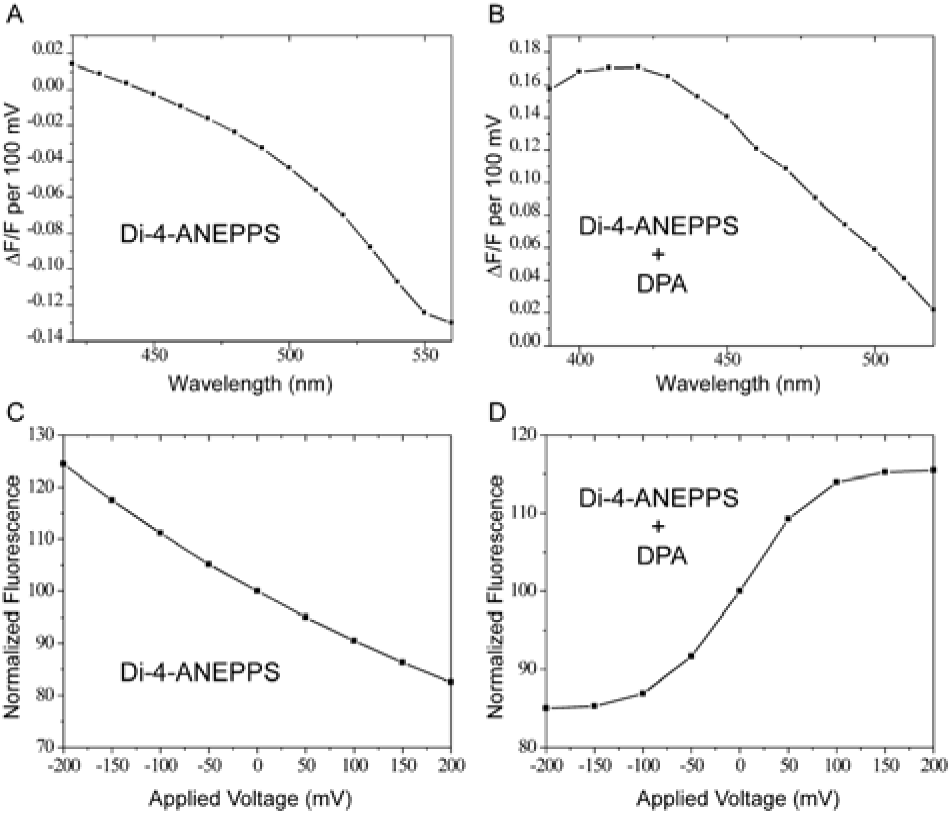
Comparison of voltage sensitivities for Di-4-ANEPPS and Di-4-ANEPPS/DPA. (A-B) Excitation wavelength dependence of voltage sensitivities for (A) Di-4-ANEPPS (long pass emission >695nm) and (B) Di-4-ANEPPS/DPA (emission > 530nm). (C-D) Voltage dependence of fluorescence for (C) Di-4-ANEPPS (550 nm excitation with >610 nm emission) and (D) Di-4-ANEPPS/DPA (450 nm excitation with >530 nm emission).

In the 2-component system, a high concentration of acceptor is required to assure that quenching will occur in the depolarized state. In practice, however, the DPA is used at concentrations below 4μM to minimize pharmacological effects. By tethering the donor and acceptor moieties, we can assure that they will always be close enough for quenching in the depolarized state and also minimize the required VSD concentration. For a quencher to move all the way across a cell membrane, a minimum 4 nm tether is needed, which is about the length of an alkyl chain with 32 carbon atoms. Since 4nm is significantly larger than the *R*_0_ of 2.6 nm for these hemicyanine/DPA donor/acceptor pairs (Fig.2), a big change of FRET efficiency, and thus the fluorescence, can be expected even though it is tethered. We decided to synthesize a series of TBFQ molecules with varying side chains and tethers, all using the aminonaphthylethenypyridinium(ANEP)-DPA donor-acceptor pair. We then determined the fluorescence responses of TBFQ VSDs to voltage steps applied to the hemispherical lipid bilayer to arrive at design principles for optimizing TBFQ voltage sensitivity.

**Scheme 1.**
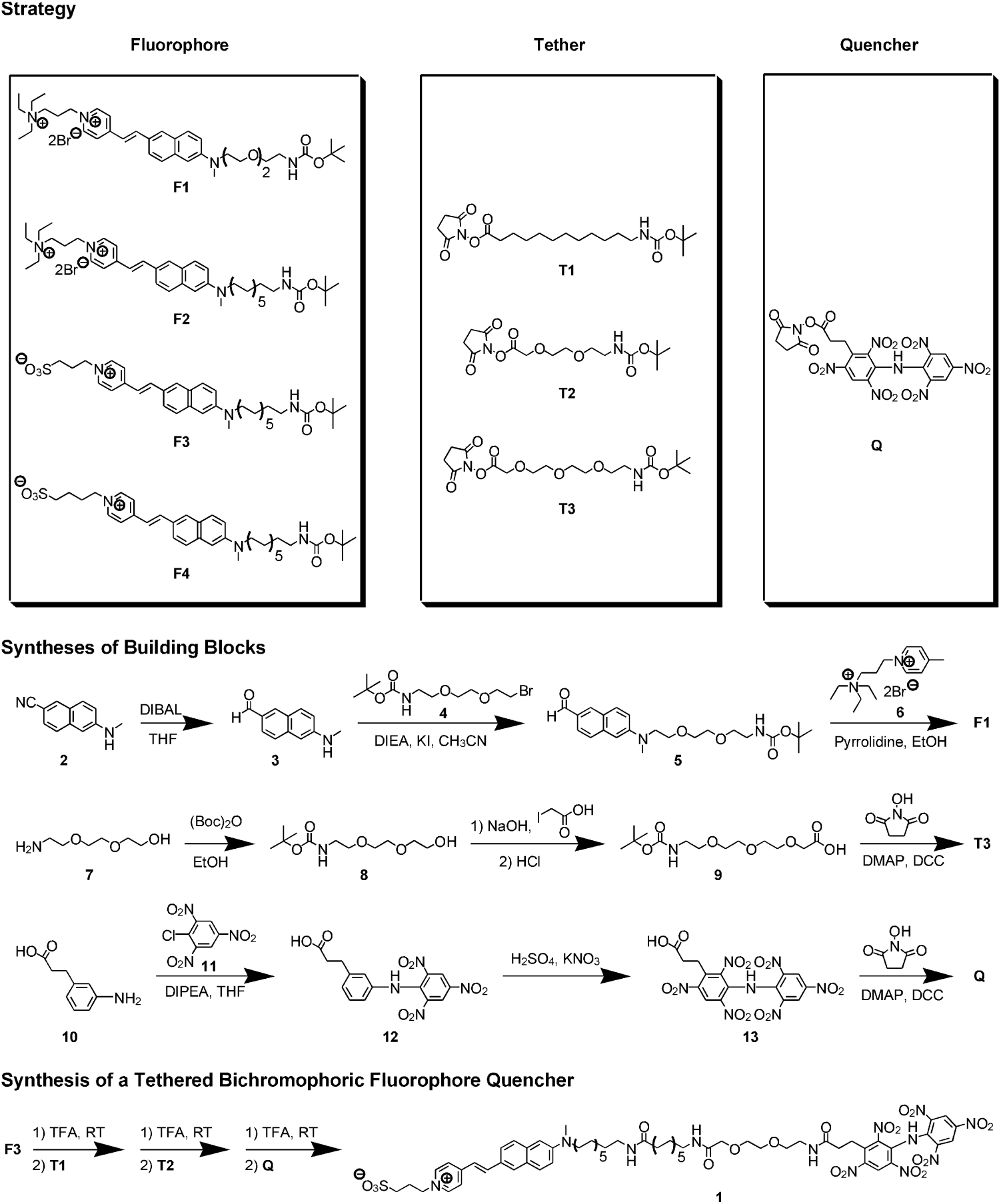
Synthesis of Tethered ANEP-DPA Voltage Sensors.

### Synthesis

A modular strategy was used to synthesize tethered ANEP-DPA dyes with tethers that are tunable both in length and hydrophilic/hydrophobic balance (see Scheme 1). The strategy relies on the convenient room temperature peptide formation chemistry between an amine group and an *N*-hydroxy succinimide ester. The building blocks consist of ANEP dyes with Boc-protected amine group (**F1**, **F2**, **F3**, **F4**), a DPA quencher with *N*-hydroxy succinimide ester group (**Q**), and various tethers with both Boc-protected amine and *N*-hydroxy succinimide ester groups at the opposite ends (**T1**, **T2**, **T3**). Hydrocarbon chain linkers **T1** serves as hydrophobic building blocks while PEG chain linkers **T2** and **T3** serve as hydrophilic blocks. A linker block can repeat or be combined with other linker blocks in any order. Synthesis of **F1**, **T3**, and **Q** are also shown in the Scheme as examples. Other building blocks, **F2**-**F4** and **T1**-**T2**, can be readily synthesized, as detailed in the Supporting Information. The last part of Scheme 1 shows synthesis of a typical TBFQ dye, **1**, which involves repetitive deprotection of an *N*-Boc group with TFA, and subsequent reaction with excess *N*-hydroxy succinimide ester at room temperature. The excess ester is used to drive the reaction to completion, but does not complicate later column chromatography purification since the product has a much higher polarity.

### Voltage sensitivity and kinetics

Table 1 shows 13 representative TBFQ dyes that have been synthesized during our quest for a superior voltage sensor. This series was guided by feedback from voltage sensitivity tests on the hemispherical lipid bilayer. The first entry, **14**, is the easiest one to prepare synthetically, but it is nonfluorescent due to the very short tether; even when stretched, this tether would be shorter than the *R*_0_ of 2.6 nm and is too short to span the thickness of a lipid bilayer. Presumably, the ANEP and DPA chromophores remain close to each other on one side of the membrane at any applied voltage. The second entry, **15**, shows a wavelength-dependent voltage sensitivity that is typical of electrochromic mechanism from the parent ANEP dyes (e.g. Di-4-ANEPPS in Fig. 3); i.e. the voltage-dependent fluorescence change displays a biphasic wavelength dependence;^5a^ a quenching mechanism is expected to be independent of excitation wavelength (e.g. the 2-component DPA/Di-4-ANEPPS pair in Fig.3). The sensitivity of electrochromic dyes is typically <20%/100mV and dye 15 is only 8%.

**Table 1.**
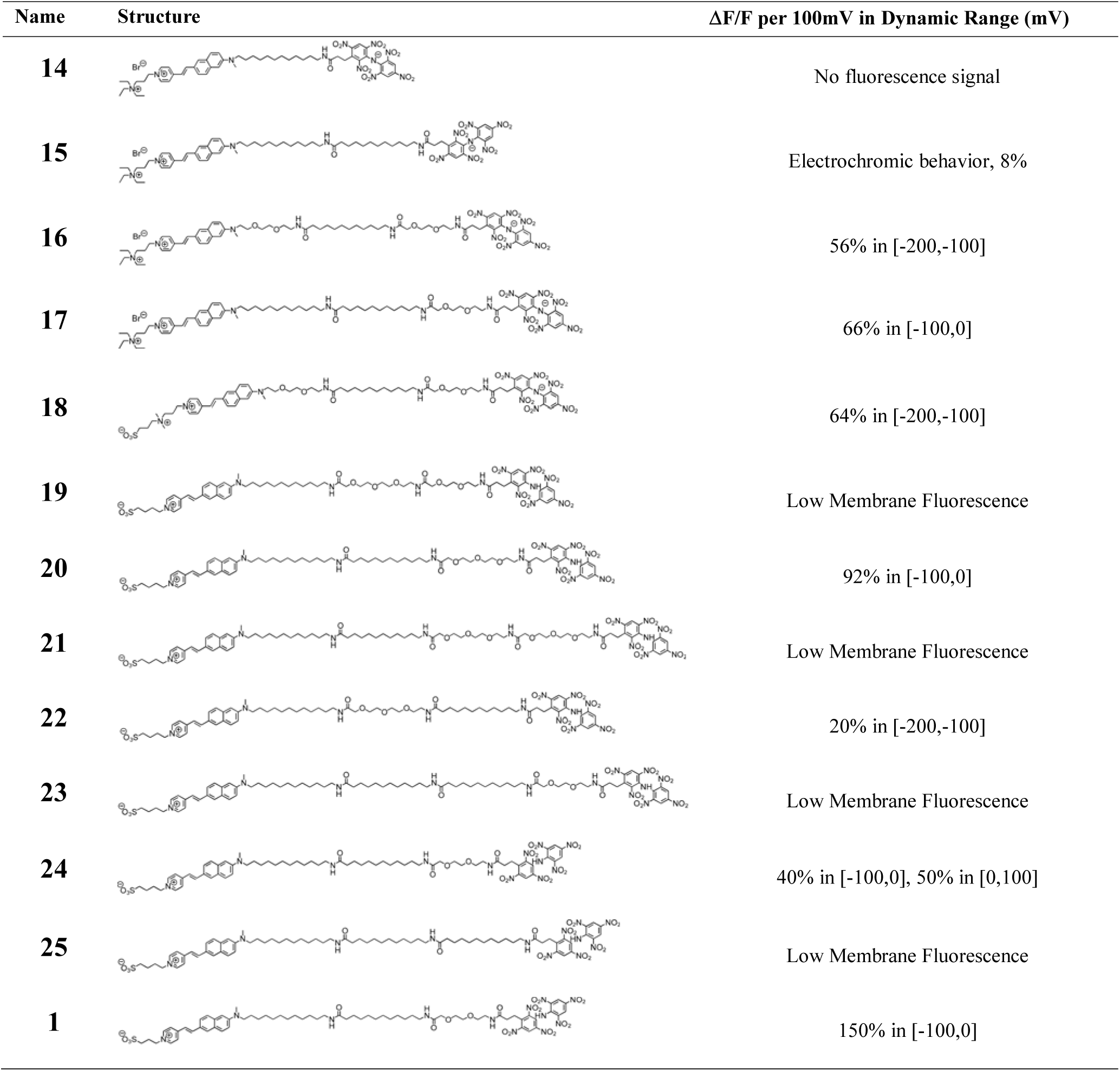
TBFQ Dye Screening

| Name | Structure | $\Delta F/F$ per 100mV in Dynamic Range (mV) |
| --- | --- | --- |
| 14 |  | No fluorescence signal |
| 15 |  | Electrochromic behavior, 8% |
| 16 |  | 56% in [-200,-100] |
| 17 |  | 66% in [-100,0] |
| 18 |  | 64% in [-200,-100] |
| 19 |  | Low Membrane Fluorescence |
| 20 |  | 92% in [-100,0] |
| 21 |  | Low Membrane Fluorescence |
| 22 |  | 20% in [-200,-100] |
| 23 |  | Low Membrane Fluorescence |
| 24 |  | 40% in [-100,0], 50% in [0,100] |
| 25 |  | Low Membrane Fluorescence |
| 1 |  | 150% in [-100,0] |

The third entry, **16**, has a longer tether at 32 atoms, sufficient to span the membrane. It also has polyethyleneglycol (PEG) moieties flanking a central hydrocarbon building block. It clearly produces high voltage sensitivity via the quenching mechanism, but the optimal sensitivity range is between −200 and −100 mV, which is outside of normal physiological range for an action potential of approximately - 70 to +30mV. A possible reason for this behavior may be that a strong hyperpolarizing potential is required to break the electrostatic interaction between positively charged side chains on the ANEP chromophore (which is also positively charged) and the negatively charged DPA. That is, it takes a large negative electric field to induce the DPA to move to the opposite side of the membrane. Another possible contribution to the stability of the folded conformation with donor and acceptor on the same side of the membrane is that the hydrophilic PEG groups are more stably solvated when they are adjacent to each other at the aqueous-lipid interface. We can distinguish between these 2 possibilities by considering compounds **17** and **18**. Both of these display better sensitivity than **16**. However, **17**, which is the same as **16**, but without the PEG moiety near the ANEP chromophore, is superior because it has its optimal sensitivity in the physiological range. In contrast, the appended negative sidechain in **18** does not appreciably shift the operating range of the voltage sensitivity displayed in **16**.

In this way, we continued to experiment with varying combinations of tethers and sidechains to optimize the properties of the TBFQ VSDs. We found that too many PEG groups (as in compounds **19** and **21**) rendered the VSDs too water soluble to become associated with the membrane, resulting in poor membrane staining. On the other hand, eliminating the PEG as in **25**, or too large a hydrocarbon region as in **23**, made the VSDs too water insoluble to be delivered to the membrane. VSDs **17, 20, 24,** and **1** all have a good balance of tether and side chain properties to produce responses in the physiological range that are larger than any previous fast VSDs.

Our best TBFQ VSD, **1**, included a single glycol ether adjacent to the DPA and a negatively charged propylsulfonate group on ANEP chromophore. It displayed a remarkable 150% ΔF/F per 100mV; that is, the fluorescence changes by 2.5 fold between −100mV and 0mV. We will now provide further characterization of the voltage dependence and kinetics of this new VSD.

### Characterization of TBFQ-VSD 1

Figure 4A shows the fluorescence signal of **TBFQ1** as a function of applied voltage. Please note that the ordinate is the total fluorescence, not ΔF/F. There is a ~2.5 fold change in fluorescence when the voltage changes from 0 to −100mV; this corresponds to the ΔF/F of 150% reported in Table 1. Sensitivity of this magnitude is unprecedented in voltage sensitive dyes. The dye was loaded inside the bubble, so a negative potential drives the quencher to the external solution in the cuvet and the fluorescence increases.

**Figure 4.**
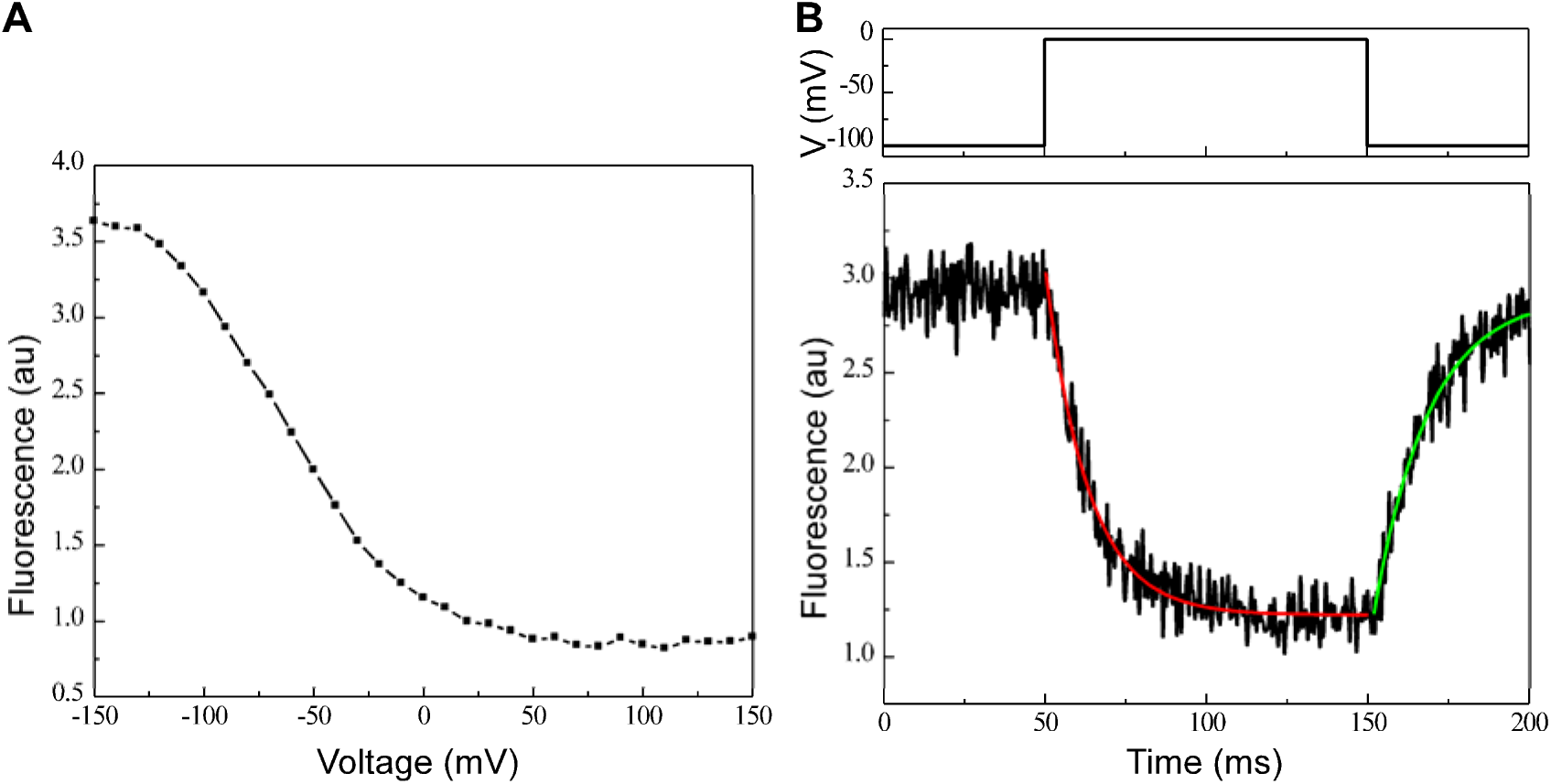
Characterization of TBFQ1. (A) Averaged (*n* = 100) fluorescence at different holding membrane potentials. (B) Kinetics of fluorescence response (bottom) to a voltage step (top). Red and green curves represent the single exponential fitting for depolarization and repolarization phases respectively. Excitation at 450nm, emission >530nm.

The kinetics of the response for **TBFQ1** is shown in Figure 2B. By fitting the curve to single exponential decay the time constant  was found to be 13 ms for the depolarization phase, and 17 ms for the repolarization phase. In literature time constants reported varies from 0.12 to 0.54 ms when DPA was used as the quencher in 2-component FRET systems.^8a, 8c^ Apparently the tether slows down the movement of quencher. However it is still on par with most protein based voltage indicators such as FlaSH,^12^ Arch (D95N),^13^ and ArcLight.^14^ It should be fast enough to detect spikes in brain or to faithfully follow cardiac action potentials in some species (including human).

In Fig. 5, we directly compared **TBFQ1** to Di-4-ANEPPS and the 2-component system of Di-4-ANEPPS/DPA using the hemispherical bilayer. Fig. 5A demonstrates the dramatic improvement in the voltage sensitivity of **1**. Figure 5B is an electrical measurement of current through the hemispherical bi-layer membrane induced by the voltage pulses. The transient currents are dues to capacitive charging of the membrane. However there is also a significant DC current for the 2-component system which could produce significant pharmacological effects *in vivo*.

**Figure 5.**
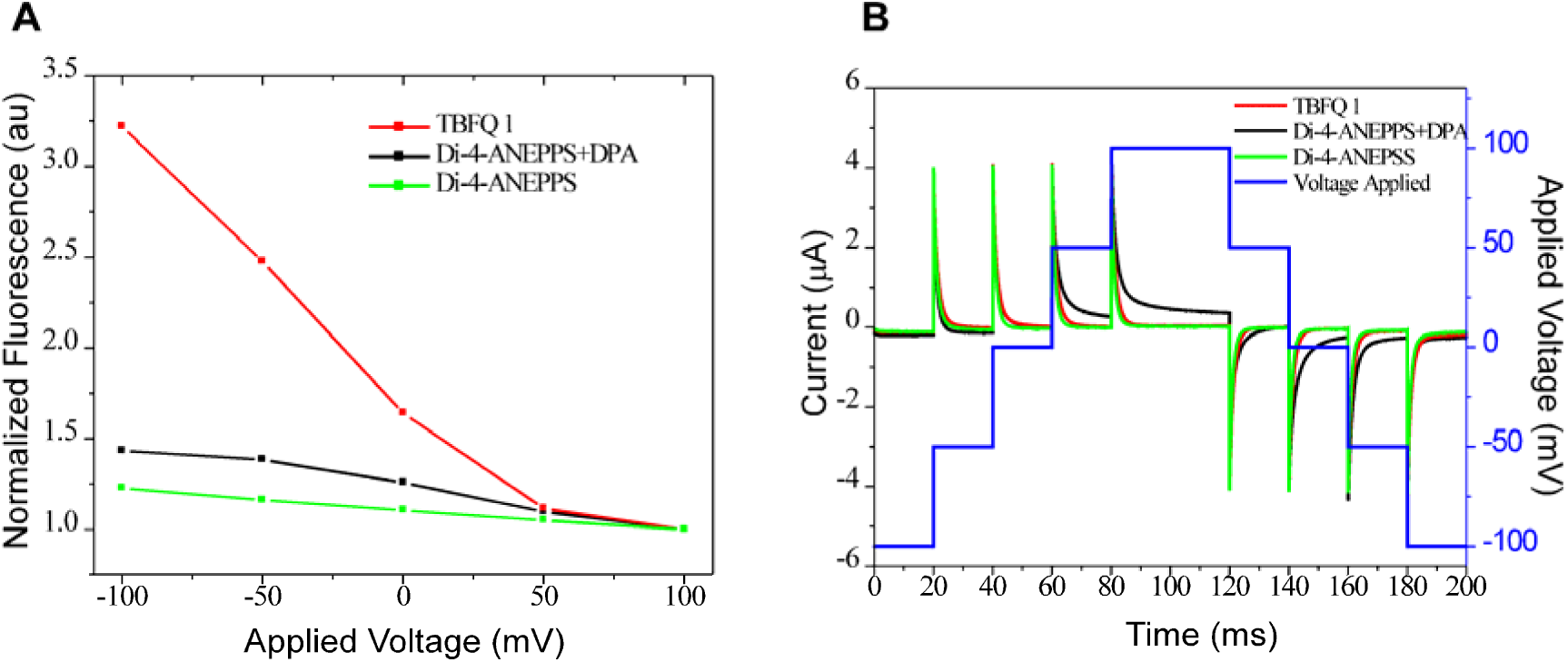
(A) Comparison of voltage sensitivity of TBFQ1, the 2-component system Di-4-ANEPPS/DPA and the electrochromic VSD Di-4-ANEPPS. (B) Current recordings in response to voltage steps for the 3 VSD systems.

The fluorescence quantum yield (FQY) for **TBFQ1** was measured to be 1.6% when bound to lipid vesicle membranes. This is approximately a factor of 15 lower than the parent ANEPPS dyes. We believe this is due to significant quenching by the DPA moiety even when **TBFQ1** is in a stretched conformation. It is clear there exists an optimal quenching efficiency for a good TBFQ dye: too little quenching will obviously not yield much voltage sensitivity; on the other hand too much quenching results in a very low baseline fluorescence signal, impeding practical applications to neuroscience and cardiology.

## Conclusion

In summary we have demonstrated a new class of tethered bichromophoric fluorophore quencher (TBFQ) voltage sensors. They have unprecedentedly high voltage sensitivities and sufficiently rapid response to detect action potentials. The fluorescence quantum yield of our best TBFQ VSD is relatively low (1.6%) compared with most organic voltage sensors. This will limit the application of this compound for fast high resolution *in-vivo* imaging of electrical activity, where low light levels will compromise the signal-to-noise ratio. However, in vitro applications, such as cell-based drug screening assays, should be significantly facilitated even with this first generation of TBFQ-VSDs, because fluorescence detection is not light intensity limited in such applications. Importantly, we have only explored a single fluorophore-quencher pair and a limited number of tethers. We are confident that improved VSDs can be developed through investigation of fluorophore-tether-quencher building block combinations.

## Associated Content

**Supporting Information**. Experimental procedures including voltage sensitivity testing apparatus setup, synthesis, characterization, ^1^H NMR and HR mass spectra for dyes. This material is available free of charge via the Internet at http://pubs.acs.org.

## Notes

The authors declare the following competing financial interest(s): The subject matter described in this article is included in patent applications filed by the University of Connecticut. LML, AC, and PY are founders and owners of Potentiometric Probes LLC, which sells voltage sensitive dyes.

## Acknowledgment

We thank Elyse Estra for technical assistance. This study was supported by the National Institutes of Health via grant Nos. EB001963.

---

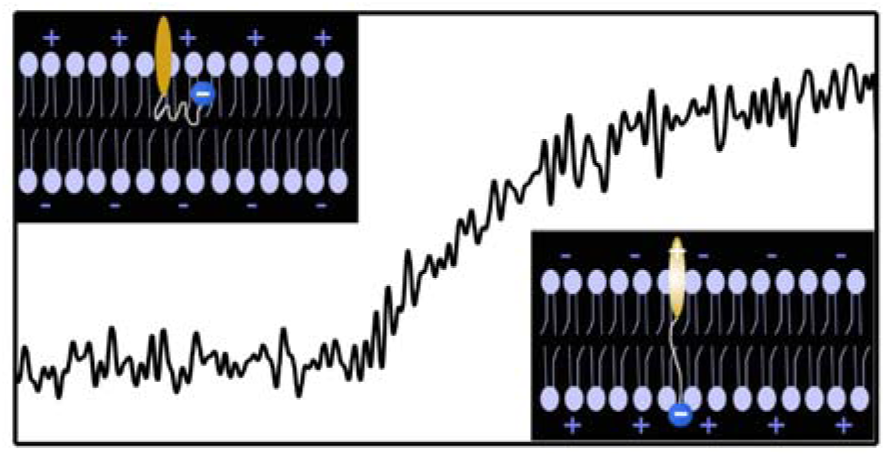
Table of Contents

